# Predicted asymmetrical effects of warming on nocturnal and diurnal ectotherms

**DOI:** 10.1101/441501

**Authors:** Marshall S. McMunn

## Abstract

Many ectotherms restrict activity to times and places with favorable temperatures. This widespread pattern of habitat use in fluctuating environments may alter predictions of how climate change will affect ectotherms. By considering time elapsed within a range of suitable temperatures as a resource, I demonstrate that warming is expected to affect thermally restricted nocturnal and diurnal activity windows asymmetrically. Under warming scenarios, thermally restricted nocturnal activity windows lengthen while diurnal activity windows contract. This divergent prediction results from the shape of the function relating time to temperature within a day, which is typically concave during the day and convex during the night. This characteristic shape is nearly universal across terrestrial environments due to the changing angle of the sun throughout each day and exponential decay of overnight temperatures. These predicted asymmetries are exacerbated by expectations of diurnally asymmetric warming (more warming during the night compared to the day). Using example data from a montane ant community, I demonstrate that, as predicted, moderate simulated warming expands activity time available to cool active species and reduces activity time available to warm active species. Together these results suggest that the time of day during which an ectotherms optimal temperature occurs can be an important factor in determining response to warming.

## Introduction

A commonly observed response to climate change within plants, animals, and fungi is to move across either space or time such that favorable abiotic conditions, including moisture and temperature, are maintained (Parmesan and Yohe 2003; Kelly and Goulden 2008; Diez et al. 2013). Historical shifts in calendar date and movement across landscapes suggest that species vary in these responses, and that particular life-history traits may increase the likelihood of experiencing negative consequences of warming (Deutsch et al. 2008). Particularly susceptible species may be long-lived and sessile, have demographic parameters sensitive to the environment (Clark et al. 2011), live in a location where climate is expected to change more quickly, have narrow thermal tolerances (Gilchrist 1995), or be engaged in tightly co-evolved mutualisms (Bennett and Moran 2015). A topic that has received comparatively less attention in evaluating capacity to adapt to temperature is activity time. Here, I aim to address how susceptibility to warming varies between organisms active at different times of the day.

Ectotherms face a daunting task safely regulating their internal body temperatures (Kearney et al. 2009). They must move through a dynamic thermal landscape in an attempt to maximize fitness through expression of a diversity of behaviors. Ectotherms often seek shelter from extreme conditions in thermal refuges, defined here as thermally buffered microhabitats, such as leaf litter, space beneath rocks, or in bodies of water. Time can only be spent in one place, and while time in a refuge may be put toward useful processes such as digestion, growth, predator avoidance (Karban et al. 2015), or colony maintenance (Gordon 1983), it comes at an opportunity cost in terms of foraging, seeking mates, defending territory, or any other engagement with the community lying outside the refuge (Amo et al. 2007). Due to movement between microhabitats, mobile ectotherm body temperature regularly deviates from local air temperature, which is often used in predictions of the fitness consequences of climate change (Gaitán-Espitia et al. 2014). The ability to decouple internal and environmental temperatures through movement and behavior confounds efforts to predict the impact of climate change on mobile ectotherms, as any one location used to characterize environmental temperature is insufficient to predict fitness outcomes.

Despite this challenge, thermal fitness curves associated with ectotherms are commonly used predict consequences of global climate change (Sinclair et al. 2016). A recent framework provides guidance as to how diurnal fluctuations in body temperature affect the ability of organisms to acclimate, and the importance of considering the effects of brief periods of exposure (Kingsolver and Woods 2016). However, many ectotherms, in particular soil-dwelling arthropods, can fully escape extreme temperature exposure, and may limit activity to times that are suitable (Sinclair et al. 2016). By using high quality thermal refuges, arthropods violate assumptions implicit in any predictions that depend on single microhabitat measures of environmental temperature. Through the use of high-quality thermal refuges, many ectotherms escape the severe consequences associated with exceeding optimal temperature (T_opt_) (Huey and Pianka 1977).

Due to the commonality of temperature based behavioral plasticity and the ubiquity of diurnal temperature variability, the duration of daily thermal activity windows could play an important role in mitigating the response of ecological communities to climate change (Kearney et al. 2009). Time elapsed within suitable temperature ranges is an important but rarely characterized resource of ectotherms. There are currently no predictions for how durations of suitable temperature may be altered for species active at different times of the day.

Daily temperature variation in terrestrial ecosystems is affected by many local factors including geography, elevation, humidity, and season (Dai et al. 1999). Despite local differences, there are several characteristics of daily temperatures that are nearly universal in terrestrial ecosystems. The sun’s local zenith results in a peak of incoming radiation at solar noon. Air temperature lags behind this peak solar radiation, as infrared radiation from the ground also contributes to warming of the air (Karl et al. 1991). Due to the continuously changing angle of the sun, a function relating time (x) to temperature (f(x)) is concave during the day. Further, temperatures surrounding solar midnight are on average, convex, as nighttime cooling results from a decelerating loss of heat that was accumulated during the day.

Global climate change is leading to uneven surface temperature warming across the globe. Poleward regions and high elevation sites are warming faster than low-lying and tropical regions. In addition to a geographic mosaic of warming, temperatures are changing unevenly in time (Davy et al. 2016). Across broad geographic regions, nighttime low temperatures are increasing faster than daytime highs. This diurnal asymmetry, represented by proportionally more warming occurring during the night, reduces the daily range of temperatures at any given site. Diurnal asymmetry of temperature may result in differential outcomes of warming for organisms depending on when during the day optimal temperatures occur.

Here, I describe and test a model predicting how diurnal temperature variation can interact with ectotherm behavioral thermoregulation based on a simple assumption that ectotherm activity is limited by local temperature and that the time elapsed within a favorable temperature range is itself a resource. I estimate the extent to which species-specific and local factors such as activity temperatures, diurnal asymmetry of warming, thermal activity breadth, daily temperature range, and day-to-day variation in temperature alter expectations of changes in available activity time under warming. Finally, I apply this approach to an observational dataset of fine-scale ant activity and temperature measurements to estimate the magnitude of these impacts on activity duration across a community of montane ants.

## Methods

I used a simulation-based approach to explore how durations of daily thermal windows are affected under a variety of warming and cooling scenarios. Simulations quantified thermal window duration across potential species thermal ranges and local environmental conditions. Each simulation resulted in a calculation of the change in the amount of time elapsed each day within a range of temperatures following warming. analyses were conducted in R version 3.5.1 (R Core Team 2017) and all figures were created using the package *ggplot2* (Wickham 2009).

### Summary of simulation methods

Activity windows were calculated as the duration of each day spent between the species minimum and maximum temperature tolerance. The key biological assumption in discussing results of these simulations as relevant to organismal fitness is that time elapsed within suitable temperature ranges itself is a limited resource for many ectotherms. This approach assumes uniformly distributed activity across suitable temperatures, but general conclusions drawn from simulations are robust to changes in activity distribution shape. I employed two common approximations of daily temperature variation, each using 10,000 observations of temperature within a day: 1) a sine wave model, oscillating between minimum and maximum daily temperatures with a period of 24 hours and 2) a truncated sine wave for daytime temperatures linked to a negative exponential function for nighttime cooling (Parton-Logan function – Eq 1 modified from (Parton and Logan 1981; Lambrechts et al. 2011).

Equation 1 – daytime temperature

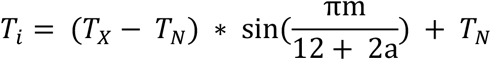

Equation 2 – nighttime temperature

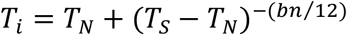

Where T_X_ and T_N_ are maximum and minimum daily temperatures respectively, T_S_ is temperature at sunset (results from Eq 1 evaluated at m = 12), m is the number of hours past sunrise, n is the number of hours past sunset, a is a lag coefficient for maximum temperature, and b is a nighttime cooling coefficient. The parameters a and b were fit in R using the nls function for data corresponding to ant activity measurements (see field methods below) and were found to provide a best fit at a = 1.15, and b = 3.37. This curve shape was then normalized such that minimum temperature = 0, and maximum temperature = 1, and scaled to the desired amplitude for each simulation.

Both functions yielded qualitatively similar results with one important exception: nighttime cooling via a negative exponential produces a long, flat tail as nighttime temperatures stabilize, exaggerating the change in activity duration for nighttime species in response to temperature shifts (Figure 1). This characteristic is realistic however, as nighttime cooling in most habitats is effectively modeled through exponential decline toward an equilibrium temperature (Parton and Logan 1981), and a Parton-Logan function was used for all simulations beyond initial comparison to a sine wave.

**Figure 1:**
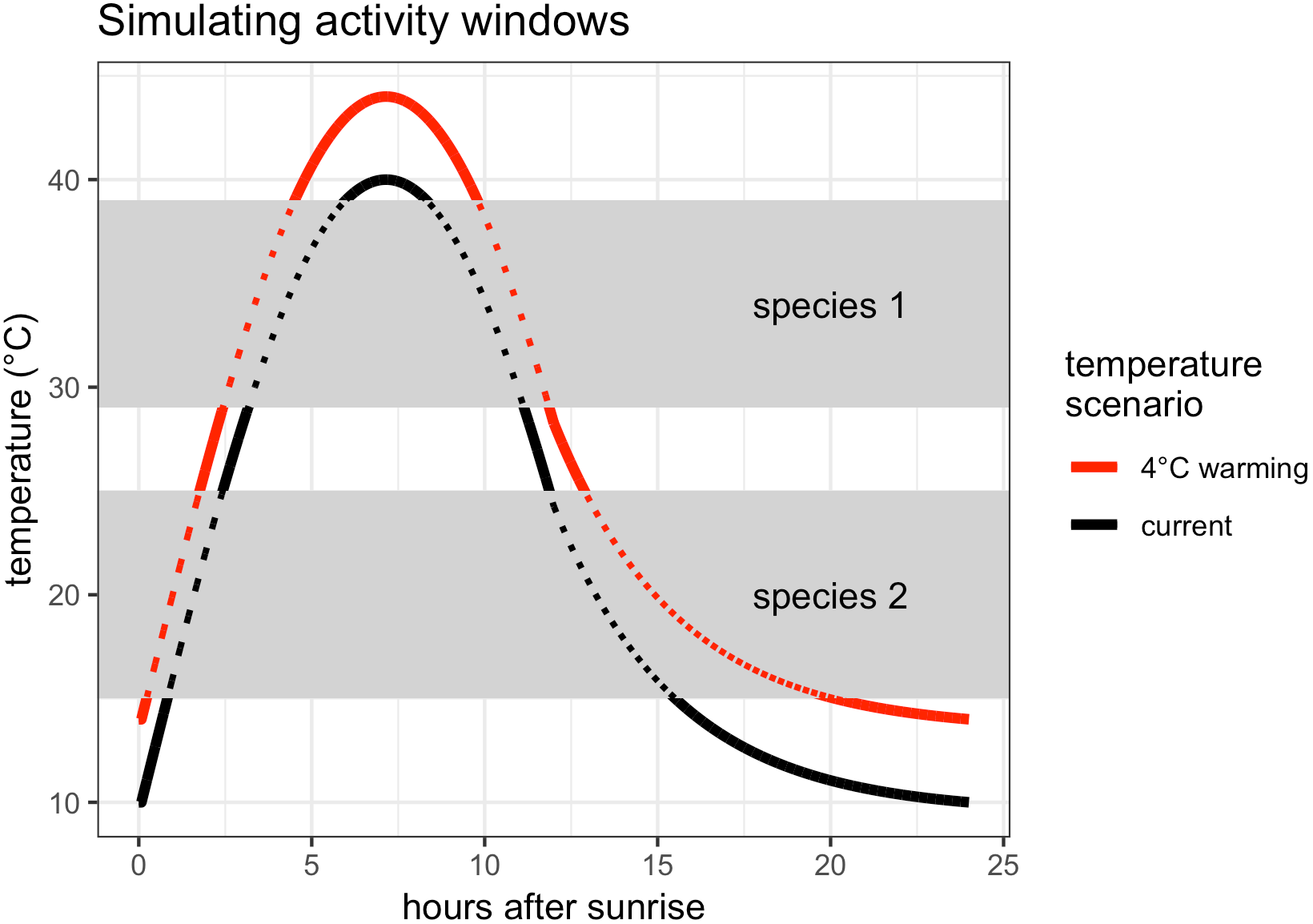
A graphical depiction of the source of divergent outcomes between a cold tolerant, morning and evening active, species and a heat tolerant afternoon active species under a warming scenario. Simulated temperature over the course of one day (black), with time measured in hours after sunrise (x-axis) and habitat temperature in °C (y-axis). The daily temperature ranges from 10°C to 40°C and follows a Parton-Logan function. A 4°C even warming scenario (increase in 4°C at all times) is also shown (red). Two potential species activity temperature windows are shown (grey), with species 1 active at warm temperatures (29°C-39°C) and species 2 active at cooler temperatures (15°C-25°C). During periods of time when the daily temperature in either scenario lies within the range of temperatures accessible to a species, the lines appear dotted, with dots evenly spaced along the x-axis. Here, species 1 (the warm active species) experiences a contraction in daily activity time available, as its preferred temperatures lie on a steeper portion of the temperature curve following warming. Conversely, following warming, species 2 (the cold active species) experiences a dramatic expansion of daily activity time available, as its activity temperature now lies on the long flat exponential decline of nighttime cooling (hours after sunrise 12-24).

All simulations were repeated with species thermal optimums (the average of thermal minimum and maximum temperatures) ranging from 10°C below the daily minimum temperature to 10°C above the daily maximum temperature (Table 1). A central set of parameters were chosen approximately in line with those observed in an empirical dataset (see field methods below) to limit exploration of parameter space to one factor at a time for clarity and understanding. These parameters were: a 10°C activity breadth for each species, a 2°C diurnally symmetric increase in temperature, a prewarming (initial) daily temperature range from 10 °C to 40 °C, and a 12-hour day and 12-hour night. These choices are not central to conclusions drawn from the models, but alignment with the empirical dataset enhances comparisons between the two.

**Table 1.** Summary of simulations performed. A summary of all simulations reported. Each row in the table represents a set of simulations across 2 changing parameters, species activity temperature and another parameter of interest. This 2-dimensional parameter space was explored using a 400×400 matrix of equally spaced values across the ranges listed above. Symbols used in the table: ΔT = the change in temperature from initial in °C, σ_s_ = breadth of species activity temperature (unless otherwise specified, σ_s_=10°C in all simulations), T_opt_ = the center of a species activity range or optimal temperature, T_x_-T_N_ = daily environmental temperature range (where Tx is maximum daily temperature and T_N_ is minimum daily temperature), SD_T_ = standard deviation of temperature at a given time resulting from day-to-day temperature variation.

| Property investigated | Temperature changes simulated | Parameter range 1 | Parameter range 2 | Figure |
| --- | --- | --- | --- | --- |
| Sine wave vs. Parton-Logan | $\Delta T = 0^{\circ}\text{C}$<br>to<br>$\Delta T = +4^{\circ}\text{C}$ | $T_{\text{opt}} = 5^{\circ}\text{C}$<br>$T_{\text{opt}} = 45^{\circ}\text{C}$ | NA | Figure 2 |
| Diurnal asymmetry of warming | $\Delta T_{\text{day}} = +2^{\circ}\text{C}$ | $T_{\text{opt}} = 5^{\circ}\text{C}$<br>$T_{\text{opt}} = 45^{\circ}\text{C}$ | $\Delta T_{\text{night}} = +2^{\circ}\text{C}$<br>$\Delta T_{\text{night}} = +6^{\circ}\text{C}$ | Figure 3 |
| Variance between days | $\Delta T = +2^{\circ}\text{C}$ | $T_{\text{opt}} = 5^{\circ}\text{C}$<br>$T_{\text{opt}} = 45^{\circ}\text{C}$ | $SD_T = 0^{\circ}\text{C}$<br>$SD_T = 12^{\circ}\text{C}$ | Figure 4 |
| Activity breadth | $\Delta T = +2^{\circ}\text{C}$ | $T_{\text{opt}} = -5^{\circ}\text{C}$<br>$T_{\text{opt}} = 60^{\circ}\text{C}$ | $\sigma_S = 0^{\circ}\text{C}$<br>$\sigma_S = 25^{\circ}\text{C}$ | Figure 5 |
| Daily temperature range | $\Delta T = +2^{\circ}\text{C}$ | $T_{\text{opt}} = 5^{\circ}\text{C}$<br>$T_{\text{opt}} = 45^{\circ}\text{C}$ | $T_X - T_N = 0$<br>$T_X - T_N = 30$ | Figure 6 |

The first set of parameters varied explored the effect of using either a sine function or a Parton-Logan function. The next scenario explored differing degrees of diurnal asymmetry of warming, ranging from even warming (+2°C day, +2°C night), to highly skewed warming (+2°C day, +6°C night). Increased nighttime warming not only modifies the minimum temperature reached, but also all temperatures between sunset and the new nighttime minimum, such that the effect of diurnal asymmetry was dampened in proportion with proximity to the temperature at sunset (Eq. 3).

Equation 3: dampening of diurnally asymmetric warming

For temperatures below *T*_*S*_:

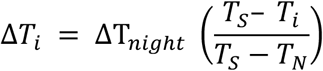

Third, the impact of day-to-day variation in temperatures was explored, by adding normally distributed noise to each daily temperature profile. This noise was added using the function *rnorm* with a mean of 0 and a standard deviation reported as a proportion of the daily temperature range (0-0.4). Next, I investigated whether species with broad thermal ranges will better tolerate environmental changes. This was accomplished by varying the breadth of temperatures available to species from 0°C to 25°C (near the extent of the daily temperature range).

Finally, I investigated the importance of daily temperature range while holding forager breadth fixed at 10°C. I simulated temperature curves ranging from 0 °C daily temperature range (unchanging temperature) to 30°C daily temperature range.

### Empirical dataset

#### Study site

I conducted field collections of ants in a small area (4.2 hectares) of mixed sagebrush shrubs and coniferous forest in the Sierra Nevada mountain range (2000m, 39.435583°N, 120.264017° W). Ants within this habitat are ground and litter dwelling, and forage on the ground, in the leaf litter, and amongst the local vegetation. The temperature on the surface of the ground, where ants frequently forage, frequently exceeds species critical thermal maximum temperatures, while nighttime temperatures are near species critical thermal minimum temperatures (McMunn, personal observations). Dominant plants include mountain sagebrush (*Artemisia tridentata ssp. vaseyana*), wyethia (*Wyethia mollis*), Jeffrey pine (*Pinus Jefferyi*), and white fir (*Abies concolor*).

#### Field Collections

I collected ground-active ants and ground surface temperature measurements using automated time-sorting pitfall traps (McMunn 2017). The traps captured ants active on the surface of the ground and within the surrounding leaf litter. Each trap contains 24 collection vials, filled with 70% ethanol, that rotate on a circular rack, with each vial being positioned under the funnel for 1 hour. I observed ants slipping quickly into the funnel and dying within seconds of submergence in the ethanol.

From a grid of 600 potential sites across the habitat, I randomly selected 127 sites for ant collection. I made collections between June 19 and October 14 2015, concentrated in one or two week sampling bouts each month. Over the season, this resulted in 3048 hour-long samples of ant abundance (24 hrs × 127 sites = 3048 hourly samples). The traps recorded temperature measurements every 5 minutes during collections using a K-type thermocouple datalogger at a height of 1-3 mm above the surface of the leaf litter or above the surface of the soil if no litter was present (McMunn 2017).

To install traps, I carefully removed leaf litter and dug a small hole approximately 20cm wide, 30cm long, and 20cm deep. I then buried the trap; replacing soil and then the leaf litter, taking care to minimize the disturbance to the surrounding litter and soil. After installation, the traps remained closed to ants for 24 hours to avoid a “digging-in” effect, when ants are initially attracted the soil disturbance following pitfall trap installation (Greenslade 1973). I separated ants from all other collected arthropods, identified each individual to species, and after confirmation of species ID’s by Phil Ward, deposited vouchers at the Bohart Museum of Entomology (UC Davis).

#### Empirical Activity Distributions

I estimated the degree of overlap between observed activity temperatures for each species and two scenarios 1) the set of all observed soil surface temperatures (from all sites, over 41,000 measurements) and 2) a simulated warming scenario (+2° C for every measure,e). Ant species thermal activity distributions were calculated using hourly temperature averages that occurred during each collection; weighted for species abundance in each sample. Empirical distribution overlap between ant species thermal activity distributions and environmental temperatures were calculated as a proportion using the R package *overlapping* version 1.5.0 (Pastore 2017). To determine the extent to which simulated warming would affect ant species active at different temperatures, I calculated empirical overlap of ant activity temperatures with 2 environmental temperature distributions: 1) all observed surface temperatures (over 41,000 measurements) and 2) even heating (+2°C for each surface temperature observation). I calculated total available activity time for each species in the 2 scenarios, observed and warming, as the proportion overlap between each species with environmental temperatures multiplied by number of minutes within a day. I then calculated the difference between these foraging windows and performed a linear regression to describe the relationship between median species activity temperature and the change in available activity time per day in response to each warming scenario (effect of warming = overlap warming – overlap observed).

## Results

I found that species activity temperatures, and when in the day those temperatures occur, are both important in determining the sign and magnitude of change in time available for activity with warming and cooling. This effect is due to the relative concavity of daily temperature curves (supplemental proof 1), which consistently have concave regions around daily maxima, and convex regions around daily minima (Figure 1).

### Warming expands activity time for cold active species

Species with activity temperatures that occur during the evening and morning, and nocturnal species, are predicted to broaden their daily temporal range due to warming (Figure 2, points 2). This broadened temporal range for species active during cooler times following warming is at first counter-intuitive, but results from a shift in timing of these colder temperatures toward relatively stable nocturnal temperature windows. This activity expansion is reversed for (likely rare) nocturnal species with activity temperatures lower than the observed daily minimum (Figure 2 – points 1). Species with optimal temperatures that lie within the concave portion of the daily temperature profile (early and late afternoon) are expected to have contracted daily activity windows following warming (Figure 2 – points 3). Again, species with optimal activity temperatures already beyond the observed daily maximum temperature will be able to take advantage of an expanded daily activity window (Figure 2 – points 4). It is reasonable to assume that activity ranges should more frequently lie within local daily temperature ranges during an organisms season of activity.

**Figure 2:**
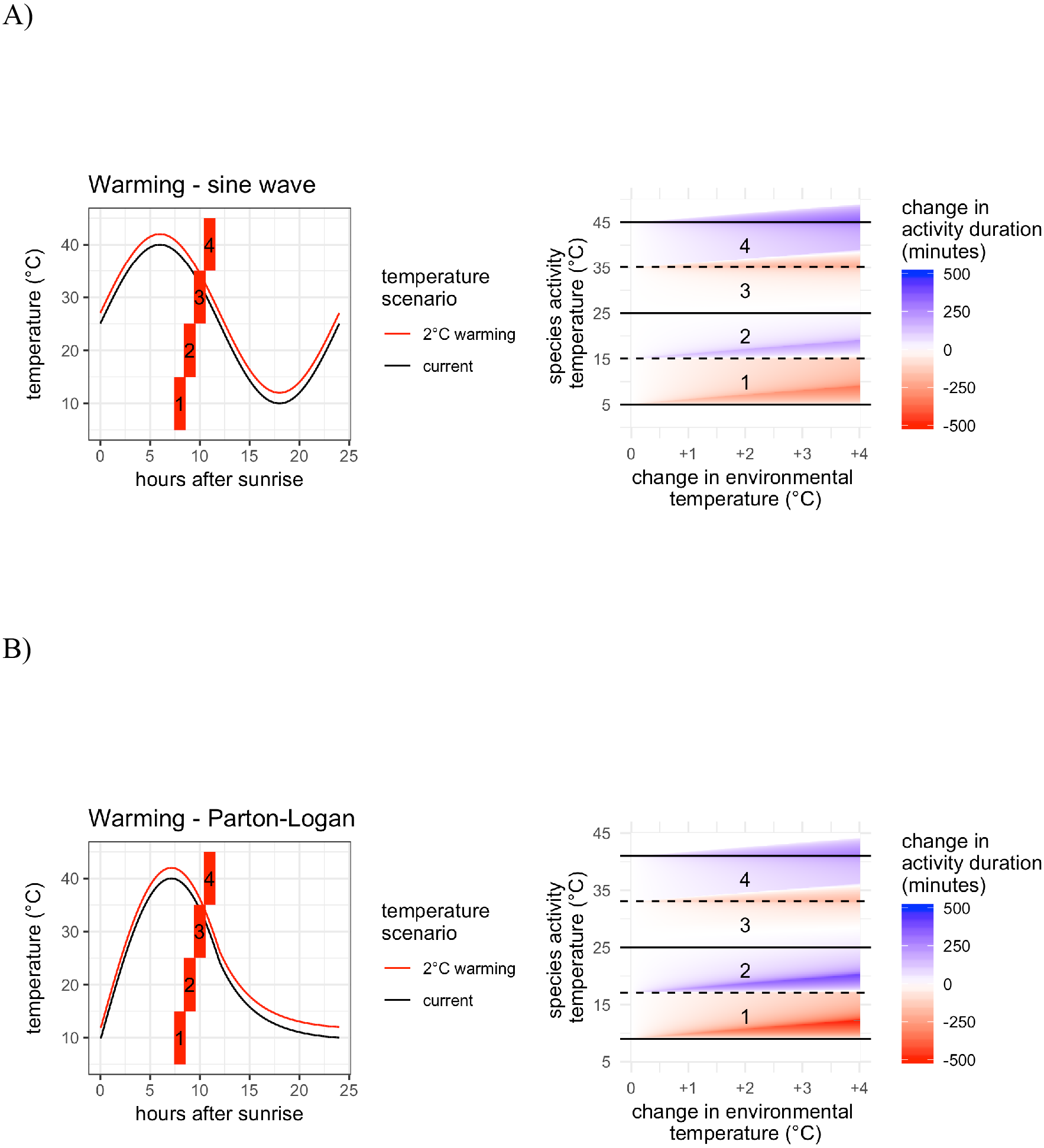
Simulations of change in available activity time with warming A) a sine wave daily temperature function and B) a Parton-Logan daily temperature function. Both panels on the left illustrate two realizations from the simulations on the right and are linked by numbers printed in each panel. On the left, colored bars represent hypotheticalspecies thermal activity ranges (each with a breadth of 10°C). The panels on the right shows the change in available activity time (minutes/day) for a simulated species with different activity temperatures (y-axis) under scenarios of temperature change ranging from 0°C to 4°C warming. The results of 160,000 simulations are plotted with white representing no change in activity duration, red as a decrease in time available, and blue as an increase in time available between the initial daily temperature profile and the new temperature profile. The magnitude of the change in activity time under any scenario is larger for cooler activity temperatures when the Parton-Logan function is used. This discrepancy is caused by the long, convex portion of the graph during the negative exponential cooling phase (hours 12-24). The regions of the plot are qualitatively describes **Species 1:** Extremely cold tolerant species have median activity temperatures below observed daily minimum temperatures and lose available activity time under warming. **Species 2:** Cold tolerant species have median activity temperatures that lie between the observed daily minimum temperature, and the daily midpoint (25 °C), and thus occupy a largely convex portion of the day. These species expand their activity time under warming scenarios, as their thermal ranges move onto a flatter part of the temperature curve (point 2). **Species 3:** Heat tolerant species have median activity temperatures that lie between the daily midpoint (25 °C) observed daily maximum temperature, and thus cover a concave portion of the graph. Under warming scenarios these species are moved onto a steeper portion of the temperature curve, reducing time available for activity. **Species 4:** Extremely heat tolerant species have median activity temperatures above observed daily maximum temperatures, and benefit from warming (point 4).

### Gradual nighttime cooling creates asymmetry in magnitude

Sine approximations of daily temperature are symmetric in their shape across their midline and spend equally as little time at the daily minimum temperature as the daily maximum temperature. This symmetry does not reflect the large amount of time spent near nighttime lows in most terrestrial habitats as radiant cooling occurs. Exponential nighttime temperature decay exacerbates differences in the expected effect for diurnal and nocturnal species (figure 2 – comparing points 3 and 4 across A and B).

### Diurnally asymmetric warming

Diurnal asymmetry of warming (more nighttime warming than daytime) increases the relative magnitude of expansions and contractions in activity time available to species active below the daily temperature midpoint. Due to this, asymmetry of warming further exacerbates the divergence of results between diurnal species and nocturnal species with warming (Figure 3 – darker colors on bottom right of right panel).

**Figure 3:**
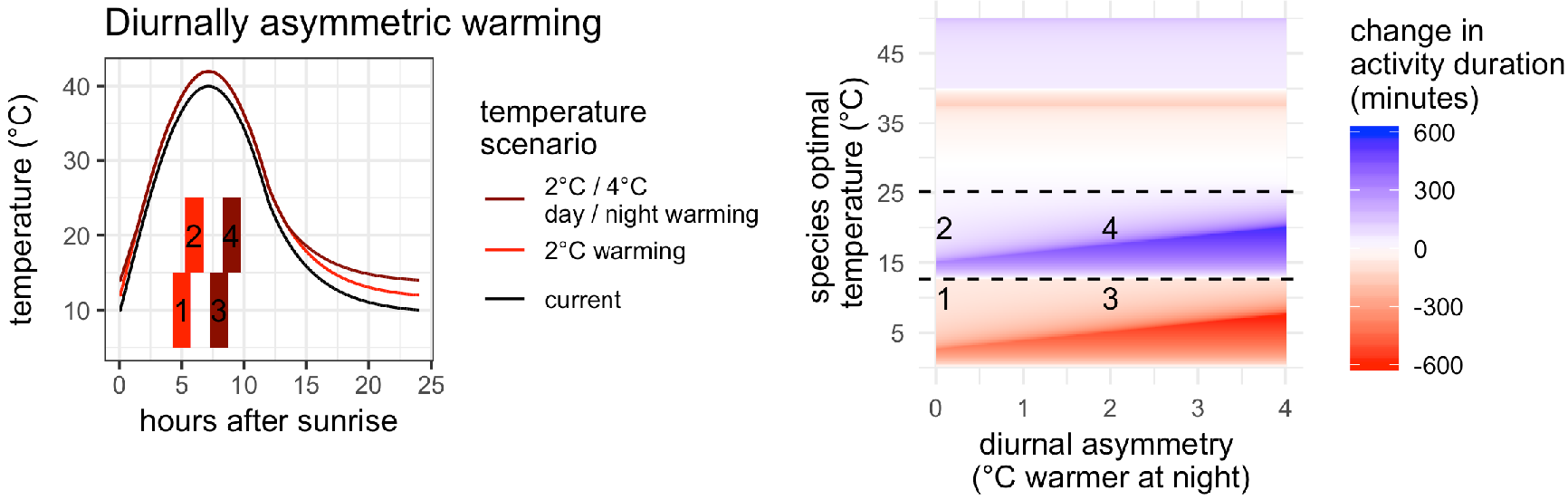
Simulations of change in available activity time for scenarios ranging from diurnally symmetric warming to highly asymmetric warming. The panel on the left illustrates four realizations from the simulations on the right, linked with numbers printed in the right panel. On the left, colored bars represent hypothetical species thermal activity ranges (each with a breadth of 10°C). The color of the bar indicates which scenario is being considered (red = diurnally symmetric warming, dark red = diurnally asymmetric warming), matching the color of the two realized temperature scenarios (red = +2°C, dark red = +2°C day/+4°C night). The panel on the right shows the change in available activity time (minutes/day) for simulated species with different activity temperatures (y-axis) under scenarios of temperature change ranging from no diurnal asymmetry to a 4°C difference in warming between night and day (+2°C day and +6°C night). The results of 160,000 simulations are plotted with white representing no change in activity duration, red as a decrease in time available, and blue as an increase in time available between the initial daily temperature profile and the new temperature profile. Diurnal asymmetry ofwarming increases the magnitude of changes in activity time for cold tolerant species (darker colors left and bottom). Diurnal asymmetry also leads to a wider range of cold tolerant species experiencing expanded activity windows (transition from blue to red at higher activity temperature).

### Variable habitats

Day-to-day temperature variation diminishes the magnitude of change in the time spent within particular thermal windows with warming (Figure 4 – increasingly weak color gradient on right side). As day-to-day variation in temperature increases, the frequency of previously rare temperatures, those occurring during fast-changing times of day, increases. Temperatures that occur during slow changing portions of the day, such as near the nighttime minimum, become relatively rarer as day-to-day variation increases.

**Figure 4:**
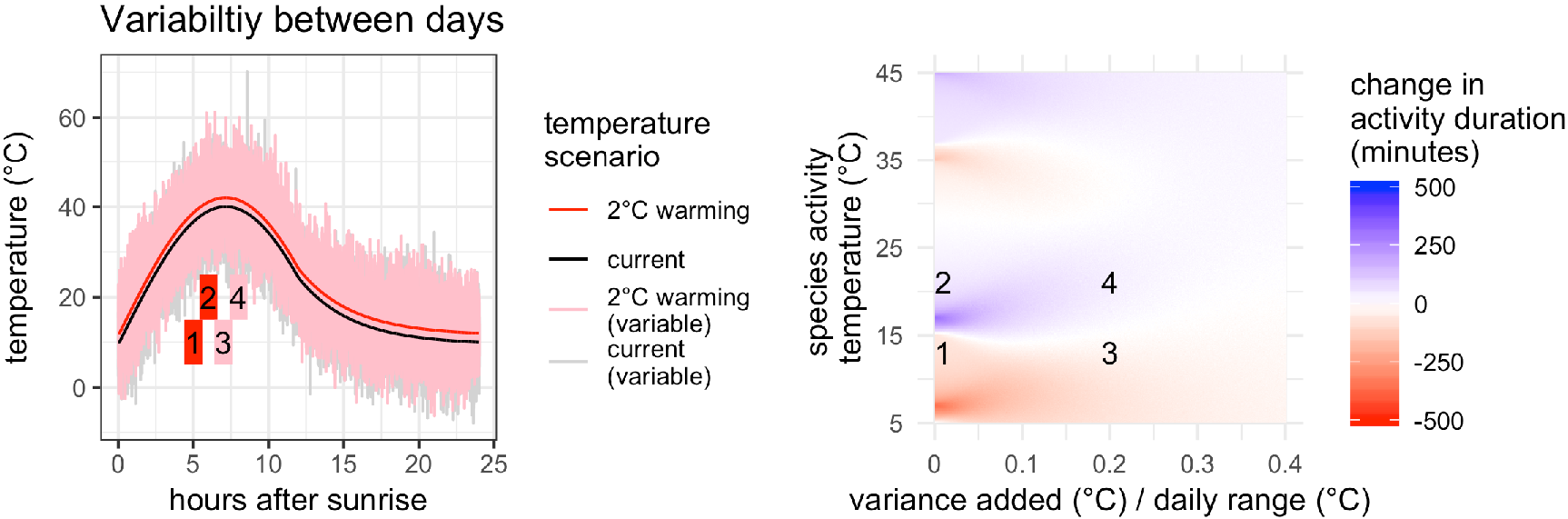
Simulations of change in available activity time for scenarios ranging from no day-to-day variation in temperature, to day-to-day variation of 40% of the daily temperature range (normally distributed with standard deviation = 12°C). The panel on the left illustrates four realizations from the simulations on the right, and is linked by numbers printed in the right panel. On the left, colored bars represent hypothetical species thermal activity ranges (each with a breadth of 10°C). The color of the bar indicates which scenario is being considered (red = warming, pink = warming with noise) matching the color of the realized temperature scenarios (black = initial, red = 2°C warming, pink = 2°C warming with random normal noise added, mean = 0, SD = 6°C). The panel on the right shows the change in available activity time (minutes/day) for simulated species with different activity temperatures (y-axis) under scenarios of temperature change ranging from no variation to large variation. The results of 160,000 simulations are plotted with white representing no change in activity duration, red as a decrease in time available, and blue as an increase in time available between the initial daily temperature profile and the new temperature profile. Day-to-day variation diminishes both the effect of activity expansion in morning/evening active species and the effect of activity contraction in afternoon active species. As variation is added around each point in time in the temperature curve, regions that are flat (e.g. nighttime) are transformed into a distribution of temperatures, rather than a narrow line. Along the length of the temperature curve, this has the effect of making rare temperatures more common, and common temperatures rarer.

### Species with broad activity temperatures

As species activity ranges become broader, the transition temperatures between activity expansion and contraction following warming move toward the daily midpoint temperature (25 °C) (Figure 5 – comparing points 3 and 4). If local species have very narrow thermal activity ranges, then most cold tolerant species would experience very weak expansion of activity windows and most heat tolerant species would experience very weak contraction of activity windows with warming (Figure 5 – right panel, left side). Conversely, if species activity ranges are very broad relative to the daily temperature range, then only species near the daily midpoint temperature experience changes in available activity time (Figure 5 – right panel, left side).

**Figure 5:**
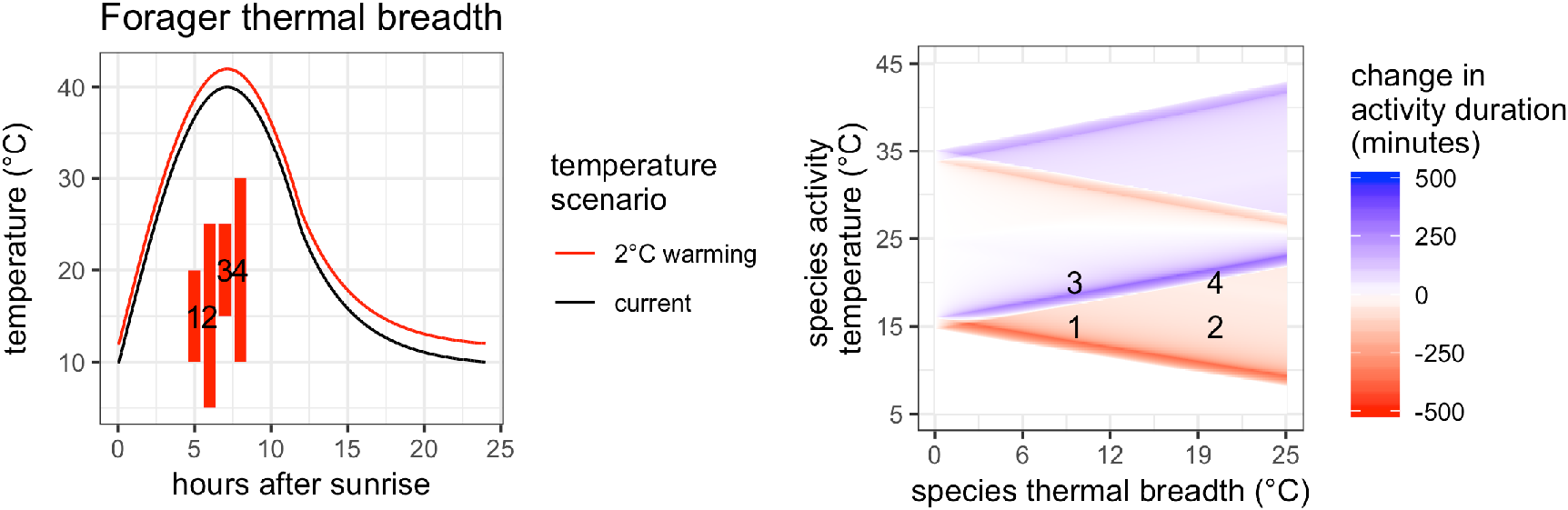
Simulations of change in available activity time for scenarios ranging from species with narrow thermal activity windows (0°C) to species with broad activity windows (25°C). The panel on the left illustrates four realizations from the simulations on the right, and is linked by numbers printed in the right panel. On the left, colored bars represent hypothetical species thermal activity ranges (two with a breadth of 10°C, and two with a breadth of 20°C). The panel on the right shows the change in available activity time (minutes/day) for simulated species with different activity temperatures (y-axis) under 2°C warming ranging from infinitely narrow ranges, to 25°C, nearly the entire daily temperature range. The results of 160,000 simulations are plotted with white representing no change in activity duration, red as a decrease in time available, and blue as an increase in time available between the initial daily temperature profile and the new temperature profile. If local species have broad thermal ranges relative to the environmental temperature range, only a small subset, with their activity windows centered at the daily temperature midpoint, experience expansion from warming due to the long cold nights (comparing points 3 and 4).

### Habitats with little diurnal change in temperature

As local daily temperature range is reduced (Figure 6 – right panel, left side) and activity breadth is held fixed, a smaller range of activity temperatures experience expansion and contraction due to concavity of the daily temperature profile. These effects, due to changes in concavity between different portions of the temperature profile, disappear entirely if daily temperature range is less than species thermal breadth (Figure 6 – 10°C thermal range).

**Figure 6:**
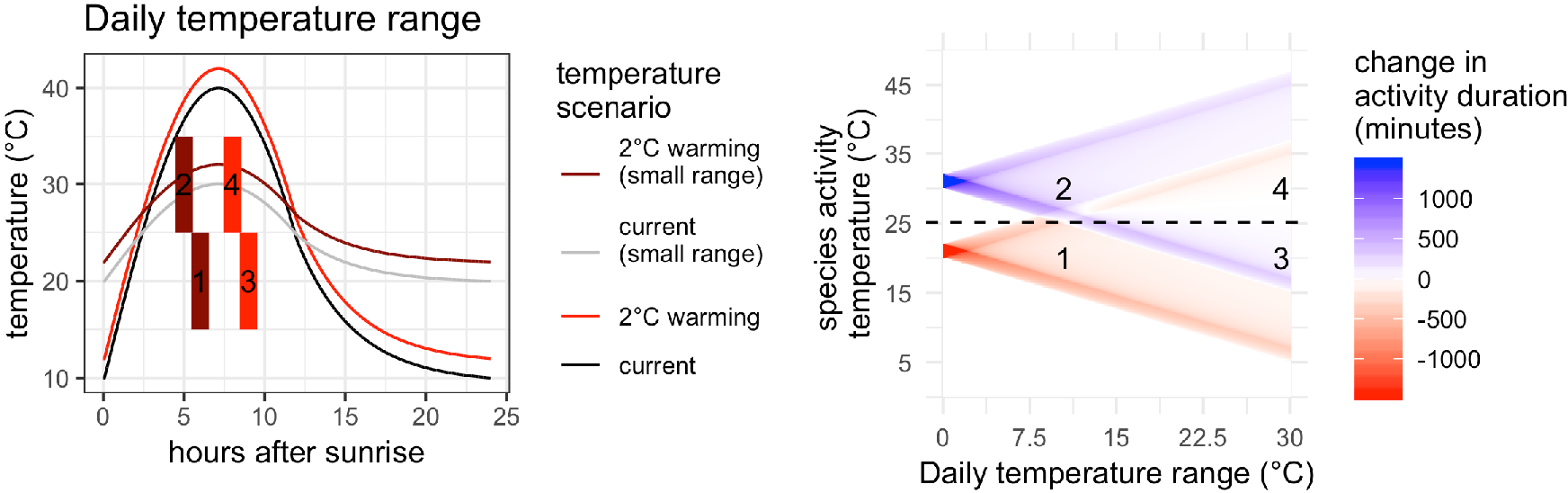
Simulations of change in available activity time for scenarios ranging from narrow daily environmental temperature range (unchanging temperature) to a daily temperature range of 30°C. The panel on the left illustrates four realizations from the simulations on the right, and is linked by numbers printed in the right panel. On the left, colored bars represent hypothetical species thermal activity ranges (all with a breadth of 10°C). The color of the bar indicates which scenario is being considered (red = warming in an environment with a 30°C temperature range, dark red = warming in an environment that has 10°C daily temperature range) matching the color of the realized temperature scenarios (black = initial, red = 2°C warming, dark red = 2°C warming, grey = initial low amplitude temperature). The panel on the right shows the change in available activity time (minutes/day) for simulated species with different activity temperatures (y-axis) under 2°C warming ranging from infinitely narrow environmental temperatures, to 30 °C daily temperature range. The results of 160,000 simulations are plotted with white representing no change in activity duration, red as a decrease in time available, and blue as an increase in time available between the initial daily temperature profile and the new temperature profile. The effect of concavity of local temperature profiles only occurs if the daily temperature range exceeds local species activity breadth (everything right of points 1 and 2). In increasingly broad daily temperature ranges, species farther from the daily temperature midpoint (25 °C) experience enhancement of activity from increased exposure to long cool nights (point 3).

### Application to observations of animal activity – ground foraging ants

In a community of montane, ground-nesting ants, species active at cooler temperatures experienced greater overlap with observed temperature following simulated environmental warming, while species active at warmer temperatures experienced activity time contraction. A linear model confirmed the negative relationship between median activity temperature and change in overlap with environmental temperature following warming (p<0.003, adjusted R^2^=0.41, t=−3.51, 15 d.f.). Changes in the duration of available activity time for each species ranged from approximately a 10-minute daily expansion to a 20-minute daily contraction of temporal range. Over 4 months (the duration of sampling in this community) 10 minutes per day accumulates to a 20-hour gain in available time.

## Discussion

The effect of warming on the availability of activity times within days is predicted to be stronger in magnitude and positive for ectotherms that are active during the temperatures that occur during the evenings, mornings, and much of the night (Figure 1 and 2). This relatively larger magnitude shift for species active below the local median temperature is enhanced due to the relative stability of nighttime temperatures during the cooling phase (negative exponential cooling - Figure 2) and diurnally asymmetric warming (relatively more warming occurring during the night - Figure 3). Conversely, animals with activity windows that lie within the range of observed afternoon temperatures will find their daily activity windows contracting with warming, as their preferred temperatures will occur during faster changing times of the day. These results are both reversed if median activity ranges lie outside observed local temperatures, a result may apply to local extremophiles or for times of the year when environmental temperature is currently unfavorable for local species.

Several aspects of local environment alter the magnitude of change in activity window size, as well as the temperatures at which sign changes in this effect occur. For an equal amount of warming, greater changes in activity time occur in more stable environments (lower daily temperature range) (Figure 6). This could lead to greater changes in available activity times in environments where high humidity or low angle sunlight limit daily temperature range, such as coastal environments, polar regions, or the tropics. Variation between days has the potential to weaken the magnitude of change in available activity times (Figure 4).

Investigating a 2°C warming scenario using ant observational data, the effect of morning activity expansion and afternoon activity contraction persisted, but at a lower magnitude than in simulations (Figure 7). This incorporates 4 months of temperature data, from mid-June to mid-October, and suggests that one very likely compensatory mechanism could be seasonal shifts in phenology. Future research characterizing seasonal phenological shifts among animals could investigate whether diurnal species demonstrate larger shifts in calendar date than species active in morning, evening, or night.

**Figure 7:**
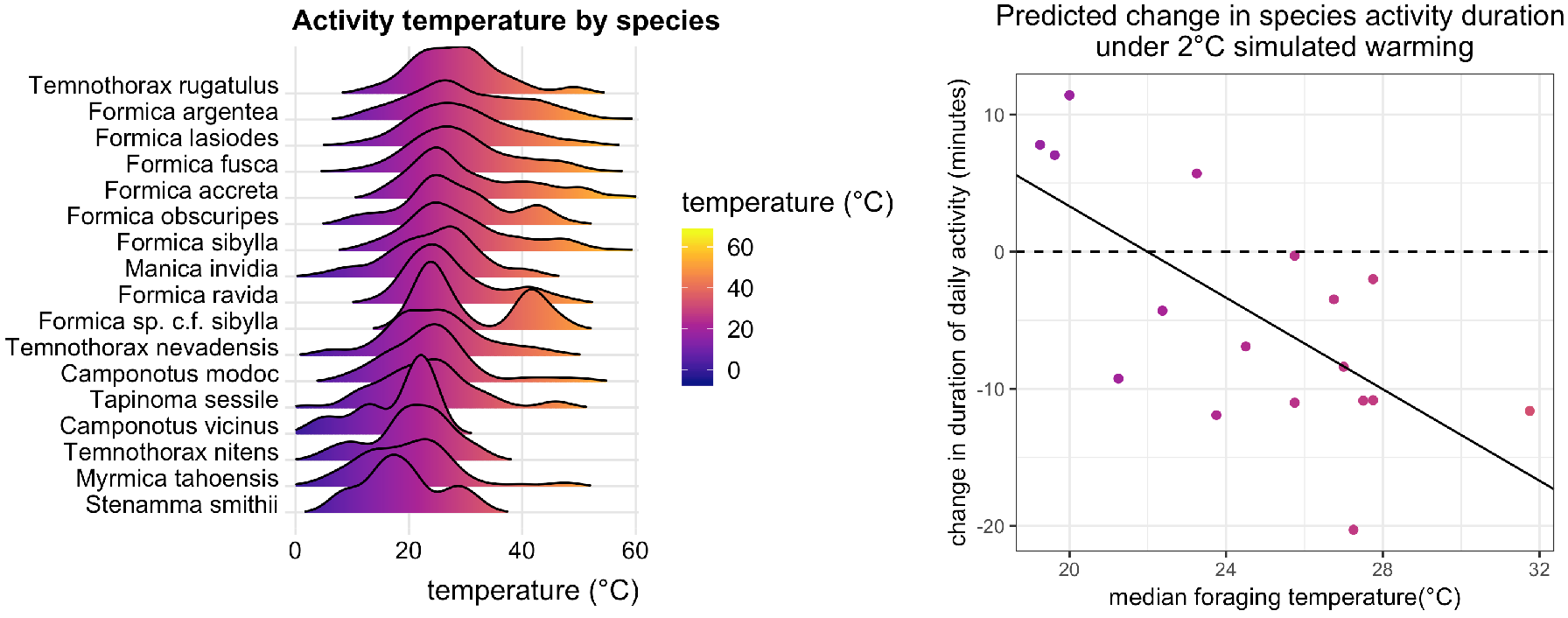
Observed median ant activity temperatures and change in estimated activity time per day with 2°C warming. Temperature measurements used in calculating empirical overlap were taken every 5 minutes, and include both day-to-day variation and site-to-site variation. Each dot is an ant species within the same community, collected at least 10 times across a period of 4 months. Available activity time for each species was calculated using an empirical estimation of overlap between environmental and species level temperature observations. A linear regression (change in activity ~ median activity temperature) is overlaid. Species with higher median activity temperatures experience a contraction in the observed environmental temperatures that overlap with their activity distributions. 4 species, all low activity temperatures, experience an increase in overlap of their activity temperatures and their environment. The sign change for change in activity time is modeled to occur at 22 °C, approximately the temperature at which the daily temperature begins a long decline toward a nighttime low (figure 1 – black line represents Parton-Logan function fitted to these temperature observations). There are no species in this community whose activity ranges are very cold or very hot.

Quality of thermal refuge in this study was assumed to be perfect, and negative consequences of temperature outside of the preferred range were not estimated. While this assumption is perhaps reasonable for ground-nesting ants, many animals are incapable of limiting exposure to such an extreme temperature. These negative effects of exceeding critical thermal maximum temperature are thought to be of particular importance, due to the steep decline in fitness once enzyme denaturation begins (Martin and Huey 2008). Even slight exposure to increased daily maximum temperatures could override the broadened activity time in the morning or evening for species with imperfect thermal refuges. Small ectotherms, and insects in particular, are notoriously hard to find if they do not return to a nest, and thus the quality of many ectotherms’ thermal refuges are not known. However, future research could compare the effect of warming on species of nesting animals with varying degrees of insulation, such as ground-nesting, tree-nesting, and litter-dwelling ants. Refuge quality could play an important role in determining whether a morning-active species benefits from an expanded available time or is harmed by exposure to now lethal afternoon temperature.

Earth’s thermal habitats are highly structured by latitude. Day length, angle of the sun, and the rate of seasonal change all shape expectations of available activity time for animals. In temperate regions, summer active animals experience a relatively short night, potentially diminishing the larger magnitude change in available activity time for morning, evening, and night-active species. Future work investigating time available within ranges of temperature could incorporate this interaction between latitude, season, and day-length.

A framework that incorporates additional factors that shape how species interact with temperature, beyond discrete availability of activity time, is necessary to accurately predict fitness outcomes of climate change for individual species of ectotherms. Ectotherm microhabitat choice, and time devoted toward thermoregulatory behaviors, such as basking, can extend activity times at a cost of decreased efficiency in foraging, hunting, mating, or any other behavior contributing to fitness. Additionally, fitness outcomes of increased temperatures depend on interactions among local species. Diel phenological mismatch could result in either enemy release or decreased resource availability depending on thermal refuge qualities among participant species.

The effect of warming on ectotherms depends on when activity times of ectotherms occur. This result is generalizable to many species that utilize high quality thermal refuges to persist in variable environments. The framework suggested here, estimating time elapsed within a range of temperature as a resource, should be included when estimating fitness impacts of climate change on species that utilize thermal refuges.

## Acknowledgements

This work could not have been conducted without my dissertation committee: Louie Yang, Jay Rosenheim, and Rick Karban. I would also like to thank Phil Ward, Brendon Boudinot, Matthew Prebus, and Marek Borowiec for their generosity and expertise in assisting with ant species identification. Thank you also to Sebastian Schreiber, who assisted in developing the supplemental proof. Finally, thanks to the insect ecology discussion group at UC Davis for helping shape me as a scientist and scholar. This project was funded in part by graduate student research grants from the UC Natural Reserve System (Mildred E. Mathias grant), the Center for Population Biology at UC Davis, and a student research grant from the American Society of Naturalists. This material is based upon work supported by the National Science Foundation Graduate Research Fellowship Program under Grant No. (DGE-1148897). Any opinions, findings, and conclusions or recommendations expressed in this material are those of the author and do not necessarily reflect the views of the National Science Foundation.

